# Scalable Production and Purification of Engineered ARRDC1-Mediated Microvesicles in a HEK293 Suspension Cell System

**DOI:** 10.1101/2024.05.14.594090

**Authors:** Kristin Luther, Ali Navaei, Leah Gens, Carson Semple, Pearl Moharil, Ilaria Passalacqua, Komal Vyas, Qiyu Wang, Shu-Lin Liu, Lucy Sun, Senthil Ramaswamy, Davide Zocco, Joseph F Nabhan

**Author notes:** **Correspondence:** Kristin Luther, Vesigen Therapeutics, 790 Memorial Drive, Cambridge, MA, 02139, USA, Joseph F Nabhan, Vesigen Therapeutics, 790 Memorial Drive, Cambridge, MA, 02139, USA. Equal contribution (Kristin Luther, Ali Navaei).

## Abstract

The engineering of human ARRDC1-mediated microvesicles (ARMMs) as new non-viral vehicles for delivery of gene therapies overcomes challenges associated with current modalities. Hurdles such as generating sufficient material to meet demand and development of appropriate characterization assays, however, persist. Our study evaluated two scalable strategies to generate GFP-loaded ARMMs, transient transfection or stable cell line-based production. The upstream ARMMs production processes utilized a suspension HEK293-derived line, termed 5B8, from Lonza. Production was evaluated in shake flasks and bioreactors. Downstream ARMMs purification processes employed Tangential Flow Filtration (TFF) and Anion Exchange Chromatography (AEX). Analytical methods included single particle analysis, ELISA, and immunoblotting. Additionally, an *in vivo* study was conducted in mice to investigate the half-life and biodistribution of ARMMs administered intravenously. 5B8 cells yielded robust production of ARMMs after transient transfection with the ARMMs loading construct or using a stable cell line containing a transgene that encodes the ARMMs loading cassette, in shake flasks or a stirred tank bioreactor, respectively. Approximately 50% of all vesicles produced were payload-containing ARMMs. ARMMs were purified by ultracentrifugation (small scale) or a combination of TFF and AEX (large scale). Both purification methods produced comparable ARMMs. *In vivo*, ARMMs showed rapid biodistribution predominantly to the spleen and liver and, to a lesser extent, kidneys, and lungs. The successful scale-up of ARMMs production illustrates the potential of engineered extracellular vesicles (EVs). Furthermore, this study highlights the potential utility of ARMMs for *in vivo* delivery of therapeutic molecules.

## 1 INTRODUCTION

Over the past 20 years, significant advances have been made in our understanding of extracellular vesicle (EV) biology, which has enabled the investigation of a range of potential applications in diagnostics and therapeutics. As naturally existing vesicles that mediate intercellular signaling, EVs possess many desirable properties for drug delivery, such as a lack of immunogenicity and toxicity [1–4], in contrast to synthetic nanoparticles [5] or viral vectors [6]. An improved understanding of the molecular composition and release mechanism of various classes of EVs has permitted the evaluation of a range of approaches for loading these vesicles with payloads that can yield specific effects in recipient cells [7].

ARRDC1-mediated microvesicles (ARMMs) are small EVs, ∼ 60 nm in diameter, that bud from the cell surface [7–9] via a mechanism driven by ARRDC1 and a number of cellular factors [10]. ARMMs have a physiological role in non-canonical NOTCH signaling [11], and were recently shown to recruit and transfer anti-oxidant proteins in response to heavy metal exposure [12]. As ARRDC1 is ubiquitously expressed, ARMMs are likely to be well tolerated, rendering this class of EVs an optimal system for the delivery of therapeutic molecules. The engineering of ARMMs to carry RNA, proteins, or Cas/gRNA ribonucleoprotein complexes as payloads for intracellular delivery has been previously reported [13]. This was achieved by engineering approaches aimed at tethering a payload of interest to ARRDC1 and overexpressing both ARRDC1 and the payloads in adherent HEK293T cells.

Although these studies provide a proof-of-concept demonstration of the possibility of engineering ARMMs to actively load cargo at the laboratory scale to support small *in vitro* and rodent studies, no evidence has been provided that ARMM production can be scaled to support clinical applications. One element that is needed for ARMMs to reach pilot scale is suspension human producer cells, because suspension cultures are easier than adherent to grow to high densities and expand in bioreactors. It is important that a human producer cell is used because the human glycosylation pattern plays an important role in protein half-life, immunogenicity, and potentially biodistribution [14,15]. Another key decision is how to introduce the transgene(s) required for production, either the flexibility of transient transfection, or the streamlined cost-effectiveness of a stable producer cell line [16]. Lastly, methods for scalable clarification and downstream processes will be required, as differential ultracentrifugation [17] becomes cumbersome with batch sizes greater than a few liters. There is no consensus on a “gold standard” method to purify EVs from any source [18,19], much less ARMMs specifically. The optimal method will likely employ at least two characteristics of ARMMs, such as their size and charge.

In this study, we evaluated both transient transfection and use of a stable cell line. We determined that Lonza’s HEK293 (5B8 cell line) [20] readily produced ARMMs which were similar to the research grade product. Next, we developed a stable producer line from the 5B8 cell line, and using this we coupled production in scalable stirred-tank bioreactors with tangential flow filtration (size-based) and anion exchange chromatography (charge-based) to purify ARMMs. Thus we demonstrated the feasibility of using this cell line to produce high-quality ARMMs at a large scale, supporting the notion that ARMMs can be engineered and manufactured to support clinical trials.

## 2 MATERIALS AND METHODS

### 2.1 Cell culture

Culture conditions: Lonza’s 5B8 cell vials were thawed for 2–3 mins in a 37 °C water bath; then, the cells were diluted in 10 mL warmed complete culture medium consisting of FreeStyle™ F17 (with 4–6 mM glutamine (and 0.2% Poloxamer 188 (cat # 24040032), and pelleted at 200 × *g* for 5 min. The cells were then resuspended in 25 mL of fresh medium. Next, 3.5–5 × 10^5^ viable cells/mL were seeded and passaged every 3–4 days in shake flasks orbiting at 120 rpm (19 mm orbit) in 5–8% CO_2_. Cryopreservation was performed in 50% CryoStor® CS10 (and 50% complete culture medium. A549 cells (ATCC, cat # CRM-CCL-185) were cultured in F12-K medium containing 10% FBS. The cells were passaged and subcultured according to the manufacturer’s recommendations.

Transient transfection: Cells were transfected at a viable cell density (VCD) of 2–2.5 × 10^6^/mL. Different ratios of DNA to transfection reagent (PEI MAX®were evaluated. DNA was 3 or 4 μg/mL of the culture volume, and the ratio was 1.375 or 1.625 to 1. The volume of the transfection mixture was 10% of the total culture volume (30 mL). VCD was evaluated daily for 3 days after transfection using an automatic cell counter, and epifluorescence microscopy images were used to confirm GFP expression.

Generation of A1-GFP stable producer cell line: Lonza 5B8 cells were transduced with a lentivirus to express a fusion protein consisting of GFP fused with ARRDC1 at its C-terminus with a short GGSSG linker. Lentivirus LVM (VB201020-1082yns)-C carrying the ARRDC1-GFP gene was generated by VectorBuilder; the titer was 6.43 × 10^8^ TU/ml. 5B8 cells were plated in 6-well plates at 4 × 10^5^ per well in DMEM containing 10% FBS and grown at 37 °C, 5% CO_2_ overnight. The medium was replaced with fresh culture medium containing 5 μg/mL polybrene before applying the virus particles to the cells with a range of MOI from 1 to 10. The medium was exchanged approximately 24 h post-infection. Forty-eight hours after the infection, puromycin was added to the media with final concentrations of 2, 4, or 6 μg/mL for selection. The cells were examined each day, and the medium was replaced with fresh puromycin-containing medium every other day. One week after puromycin selection, the cells were transferred to T75 flasks containing puromycin-free medium. Once the cells reached 80–90% confluency, they were transferred into F125 shaker flasks and grown in suspension in Lonza complete culture medium (FBS-free).

Cell culture expansion in 1 L shake flasks: The stable cells were expanded in 1 L shake flasks after seeding at 5 × 10^5^ or 1 × 110^6^/mL. They were grown in a 300 mL working volume per flask for 8 days, with samples taken each day to check cell density and viability. The 1 L shake flask seed train for bioreactor inoculation was cultured for 4 days.

Bioreactor culture: The 3 L bioreactor (BioBLU® 3c Single-Use Bioreactor, cat # 1386121000) was equilibrated and seeded at a VCD of 5 × 10^5^ cells/mL, in a total working volume of 3 L. Batch culture was conducted for 6 days. The tank was stirred at 100 rpm, and the following process setpoints were maintained: pH 7, dissolved oxygen 50%, and temperature 37 °C. These parameters were maintained automatically using a G3Lab Bioprocess Controller.

### 2.2 Isolation of EVs

Cell and debris removal: Raw conditioned medium (CM) was collected, cells were pelleted by centrifugation at 300 × *g* for 10 min, and large debris was pelleted by centrifugation at 10,000 × *g* for 30 min. This 2-step centrifugation procedure was used as a pre-measurement sample preparation step for raw CM that did not undergo downstream purification.

Ultracentrifugation (UC) purification: The UC was performed in a Beckman Coulter Optima XE at 4 °C for 2 h at 174,000 × *g* in a swinging bucket rotor (SW 32 Ti).

Downstream purification processing: After 6 days in the bioreactor, endonuclease was added to the bioreactor to a final concentration of 50 U/mL of the cell culture, and host cell DNA was digested for 2 h at 37 °C under agitation. Then, a predefined concentration of NaCl solution was added to the bioreactor to stop the nuclease reaction and reduce aggregation. Cells and cell debris were filtered out by depth filtration, followed by 0.2 µm bioburden reduction filter. The filtrate after depth and sterile filtration is called clarified CM. The clarified CM was then concentrated and diafiltered by tangential flow filtration (TFF) using hollow fiber filters with a molecular weight cut-off of 500 kDa. During diafiltration, the EVs were buffer-exchanged into a Tris-based buffer at pH 7 with a final NaCl concentration of 100 mM. The concentrate was further purified by anion-exchange chromatography using a monolithic column and polished by multimodal chromatography on an ÄKTA pure™ 150 M. The EV eluate was collected by pooling the fractions containing the highest intact EV concentrations. The final purified EVs were concentrated and diafiltered in PBS using TFF, followed by a final 0.2 µm sterile filtration.

### 2.3 Characterization of EVs

Nano particle tracking analysis (NTA): The samples were diluted in reverse osmosis distilled (RO/DI) water to within the linear working range of the NTA instrument (PMX 120 ZetaView® Mono Laser, Particle Metrix), with 50–200 particles detected on the screen. All samples were analyzed using the same procedure, with a sensitivity of 82.4, a shutter of 100, and two videos captured at 11 positions with a high frame rate.

Protein extraction from cell lysates: In total, 1–2 × 10^6^ cells were lysed in NP-40 lysis buffer containing a protease inhibitor cocktail. The cells were pelleted and frozen at −80 °C at Lonza before shipping on dry ice to Vesigen. Pellets were thawed on ice and resuspended in 50 µL PBS before adding 200 µL NP-40 lysis buffer. The lysate was vortexed and incubated on ice for 20 min before centrifugation at 16,000 × *g* for 10 min at 4 °C. The protein concentration was determined using the Pierce™ Rapid Gold BCA Protein Assay Kit according to the manufacturer’s instructions.

Western Blot: For cell lysates, 25 μg of protein were run per lane. For EVs, a consistent number of particles, 1 × 10^8^, were run per lane under reducing conditions to blot for ARRDC1, GFP, and Calnexin or under non-reducing conditions to blot for tetraspanins (CD9, 81, and 63). The EVs were lysed in 4x Laemmli buffer, heated to 95 °C for 10 min, and then run on a gradient gel at 200 mV for 20–25 min. Proteins were transferred to nitrocellulose membranes and blocked in 5% milk/TBS-T for 1 h at room temperature. Membranes were incubated at 4 °C overnight with primary antibodies to ARRDC1, GAPDH, Calnexin, GFP, CD9, CD63, or CD81. Clones, sources, and dilution factors are listed in Table 1.

**TABLE 1.**
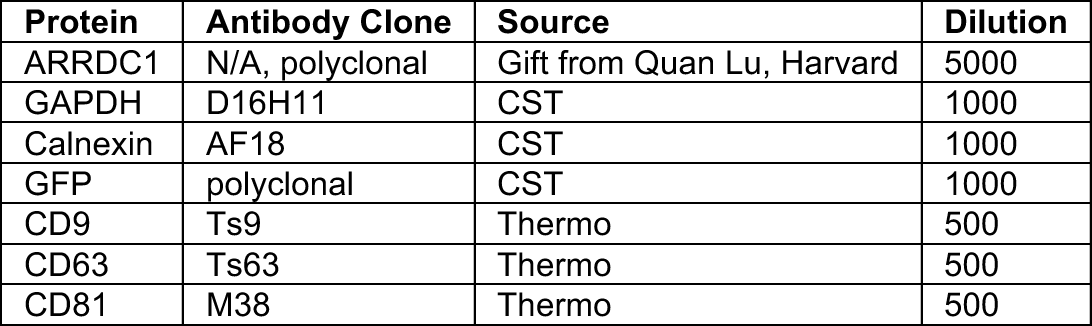
Antibodies used for Western blotting.

| Protein | Antibody Clone | Source | Dilution |
| --- | --- | --- | --- |
| ARRDC1 | N/A, polyclonal | Gift from Quan Lu, Harvard | 5000 |
| GAPDH | D16H11 | CST | 1000 |
| Calnexin | AF18 | CST | 1000 |
| GFP | polyclonal | CST | 1000 |
| CD9 | Ts9 | Thermo | 500 |
| CD63 | Ts63 | Thermo | 500 |
| CD81 | M38 | Thermo | 500 |

Protein assay: conditioned media, UC supernatant, UC pellet, and samples from each step of the downstream process were analyzed using the Bradford assay (Thermo cat # PI23238) in a 96-well format. For samples with protein concentration less than 120 μg/mL, the micro-Bradford 96-well plate format was used, according to the manufacturer’s instructions.

dsDNA was measured using a Quant-iT™ dsDNA Assay Kits according to the manufacturer’s instructions.

Single particle analysis: EVs in CM were labeled with CellTrace™ Far Red Cell dye, and unincorporated dye was washed out by size-exclusion chromatography (SEC) as previously described [21]. Data analysis and reporting followed the recent guidelines for extracellular vesicle flow cytometry experiments [22].

Uptake and GFP ELISA: A total of 1 × 10^4^ A549 cells per well were seeded and allowed to expand to approximately 70% confluence for 24 h. The medium was replaced with fresh growth medium containing appropriate concentrations of ARMMs to obtain 1 × 10^4^ and 1 × 10^5^ particles per cell (assuming 2 × 10^4^ cells/well) in triplicate. The cells were then incubated with ARMMs for 24 h. To ensure a complete wash-out of non-vesicular GFP, the wells were rinsed three times with media and once with PBS and then lysed in presence of an extraction enhancer. The lysates were assayed undiluted. The GFP concentrations in the lysates were compared using a 2-way ANOVA with Šídák’s correction for multiple comparisons. A P value of ≤0.05 was considered significant.

### 2.4 Biodistribution and pharmacokinetics

C57BL6 mice were sourced from and housed at the Charles River Accelerator and Development Lab (CRADL®) vivarium. They were maintained on a 12 hour light/dark cycle, and provided with food and water ad libitum.

EVs were injected via the retro-orbital route, 0.2 mL per mouse at a concentration of 3.3 × 10^12^/mL. There were two groups of four mice, and a sham group, which received 0.2 mL of PBS vehicle. The first group had blood collected by cheek bleed 10 min post-injection, and the mice were sacrificed 2 h post-injection for tissue collection, including a second blood draw. The second group underwent blood collection at 30 min post-injection, and the mice were sacrificed at 6 h post-injection for tissue collection, including a second blood draw. This resulted in four time points for plasma GFP concentration and two time points for tissue biodistribution. To obtain plasma, blood was collected in ethylenediaminetetraacetic acid-coated tubes, and the cells were pelleted at 3,300 × *g*. Prior to tissue collection, the mice were perfused with PBS. Tissues were snap-frozen and homogenized using HG-400 MiniG® homogenizer. Protein was extracted and the same amount of total protein (300 μg) was used as the input for each well in a final volume of 50 µL. Plasma was assayed at a 1:5 to 1:100 dilution in a final volume of 50 µL per well. All dilutions were prepared in the extraction buffer of the ELISA kit to ensure EV lysis.

To determine the total amount of GFP in the plasma, the concentration obtained from ELISA was multiplied by the total blood volume of the mice, assuming a blood volume of 72 mL/kg mouse body weight [23]. Circulating half-life was determined using an exponential one phase decay model with least squares regression. The Shapiro-Wilk test showed that the data points were normally distributed. The 95% confidence interval of the half-life was calculated, and the R^2^ value was 0.999.

## 3 RESULTS

### 3.1 5B8 cells in suspension culture readily produce ARMMs

The transient transfection of ARRDC1-overexpressing plasmids into cells drives ARMM production *in vitro* [10]. However, the transfection parameters and yield may vary depending on the producer cell line. Here, we used an HEK293-derived cell line, 5B8, which has previously been used for the production of cell- and gene-therapy products with scalable manufacturing processes [20]. Four different transient transfection conditions were evaluated (G1-4). We varied the amount of plasmid DNA and the ratio of PEI to DNA (Figure 1A) and examined their effects on cell viability, ARMM loading, and ARMM production. Expectedly, using PEI reduced cell viability compared to that of the control; however, the viability remained above 80% until the harvesting of CM on day 3 post-transfection (Figure 1B, C). Similar levels of ARRDC1-GFP (A1-GFP) expression were observed in 5B8 cells compared to other HEK293-derived cell lines. Increasing the DNA concentration or PEI:DNA ratio did not pronouncedly affect the viability or expression of A1-GFP (Figure 1D, E).

**FIGURE 1:**
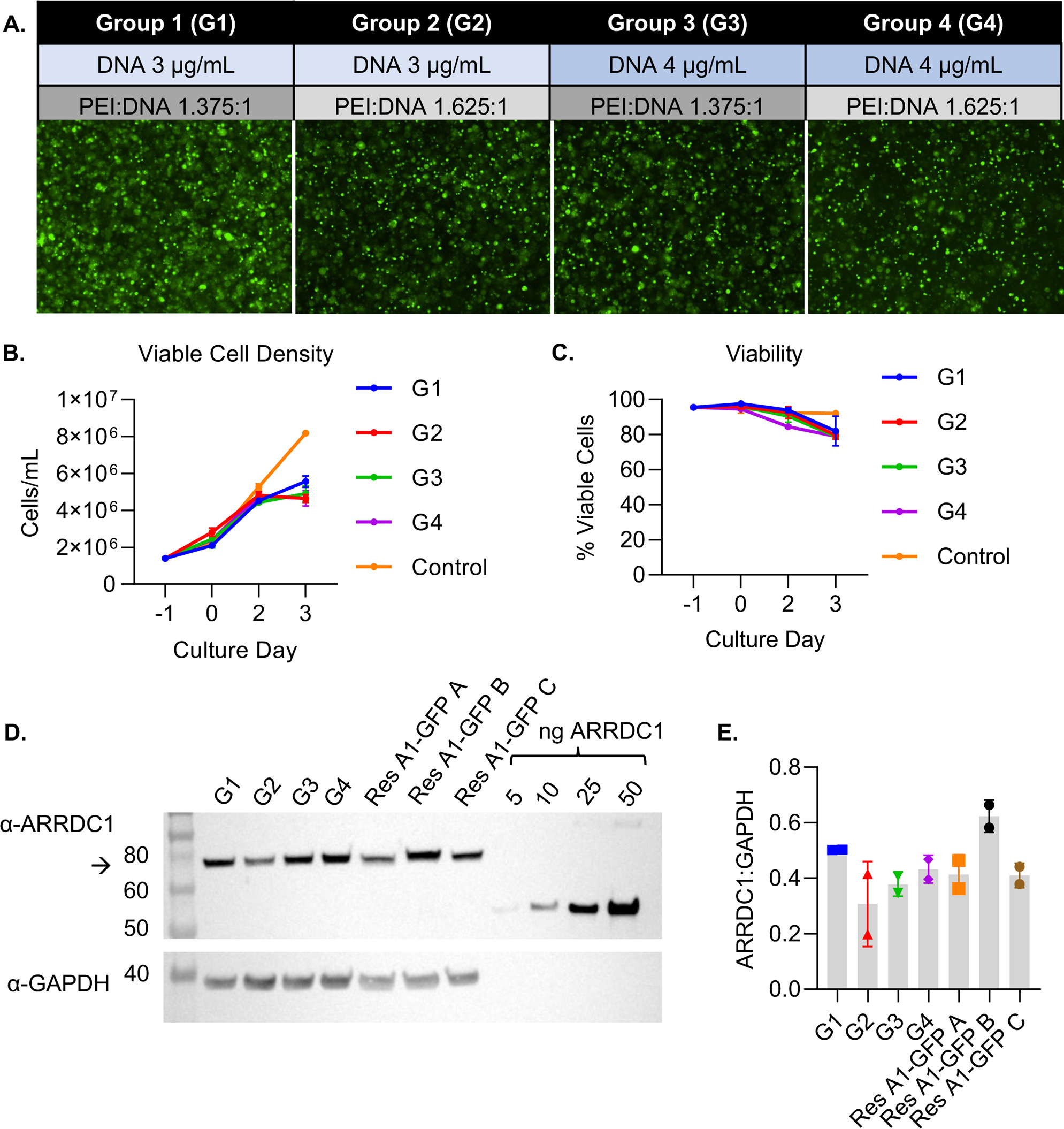
Transient transfection and expression of ARRDC1-GFP in cell lysates. **(A)** Schematic of transfection conditions and fluorescent images of the cells in each group of two shake flasks. **(B)** Viable cell density (VCD) at seeding (−1), transfection (0), and days 2 and 3 post-transfection. **(C)** The percentage of viable cells as assessed by the NC-200 cell counter. **(D)** Representative Western blot image of the cell lysates from groups G1–4 and three research HEK293 cell lines expressing A1-GFP (Res A1-GFP A, B, and C). Recombinant ARRDC1 standards are included at 5, 10, 25, and 50 ng/lane. The arrow indicates the ARRDC1-GFP fusion protein with the expected molecular weight of 73 kDa. **(E)** A1-GFP band pixel densities normalized to GAPDH (n = 2).

The particle concentrations were determined using nanoparticle tracking analysis (NTA). Similar numbers of particles were observed in G1–4, ranging from 4.7–6.3 × 10^10^/mL vs. 2.2 × 10^10^/mL in control CM, a ∼2–3-fold increase in particle concentration compared to the control CM (Figure 2A). Induction of particle production by ARRDC1 overexpression has been previously described [13]. Consistent with the smaller size of ARMMs, we also observed that the particles derived from ARRDC1-overexpressing cells had a higher proportion of smaller particles than the control (Figure 2B).

**FIGURE 2:**
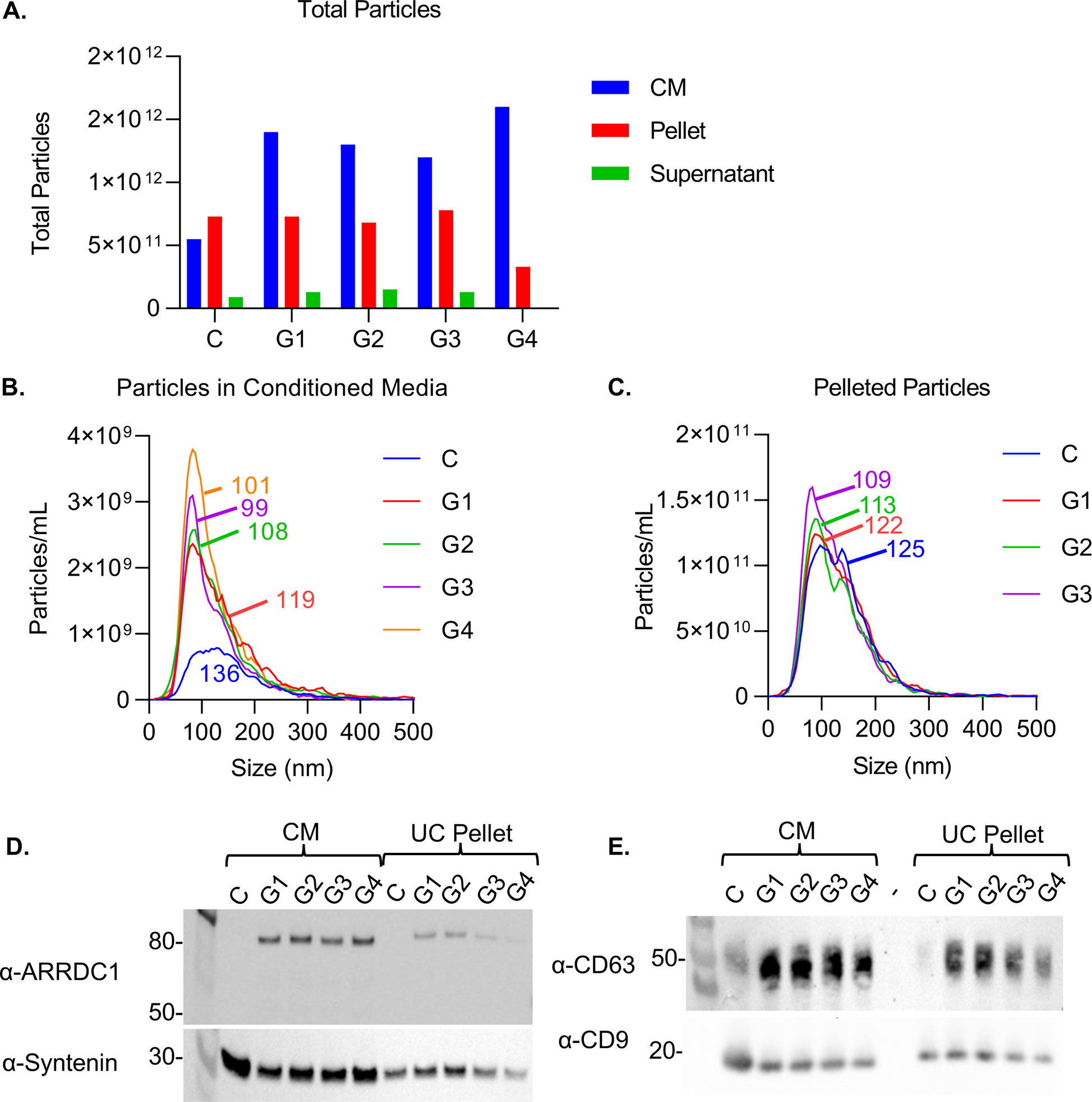
Characterization of ARMMs purified by ultracentrifugation. **(A)** Particle concentration was determined by NTA, and the mass balance of particles before and after ultracentrifugation was calculated. **(B–C)** The size distributions for particles in the CM and resuspended pellet were also determined by NTA, and the median size was labeled. **(D)** Western blot for ARRDC1 and the EV marker protein Syntenin in the CM and UC pellet at 4.5 × 10^8^ particles/lane. **(E)** Western blot for CD63 and CD9 in the CM and UC pellet at 4.5 × 10^8^ particles/lane.

Ultracentrifugation was conducted on 25 mL of CM after removal of cells and cell debris from each batch, and pellets were resuspended in 250 µL of PBS, representing a 100x concentration factor. G4 samples were excluded from the analysis. The trend toward a smaller median particle size with ARRDC1 overexpression noted in the CM was maintained in the EV pellets (Figure 2C).

Particles from the CM and EV pellets were subjected to Western blotting (Figure 2D, E). 4.5 × 10^8^ particles were loaded per lane. As expected, ARRDC1 was not detectable in the control particles after a short exposure time, and its amount in the CM was similar for all four transfection conditions, suggesting that ARRDC1 overexpression robustly increased the generation and loading of ARMMs, whereas the steady-state levels of ARMMs produced by 5B8 cells were relatively low. CM from transfected cells had higher A1-GFP levels than the UC pellet. G1 and G2 had higher A1-GFP levels in the UC pellet than G3 and G4, suggesting that 3 μg/mL of plasmid of DNA is more appropriate for production than 4 μg/mL.

All of the pelleted EVs had lower ARRDC1 levels than in the CM, suggesting that some ARMMs may have ruptured during UC or CM was contaminated with non-vesicular A1-GFP. Syntenin, which is commonly used as an EV marker [24,25], was present in all samples. We observed higher levels of syntenin in control CM, suggesting that syntenin may be less enriched in ARMMs. We further evaluated the levels of known EV markers, CD9 and CD63. CD63 was enriched in EVs derived from ARRDC1-overexpressing 5B8 cells compared to that in the control. Notably, this is the first demonstration of ARMM production by a suspension cell line cultured in a fully defined serum-free medium.

Samples from the CM, UC pellet, and supernatant were further evaluated using an anti-GFP ELISA. The signal from the total amount of GFP in the UC pellet was lower than that in the CM (Supplemental Figure 1), which was consistent with the immunoblot analysis (Figure 2D, E), suggesting that UC may compromise the particle integrity or that the CM was contaminated with cell lysates.

To determine the percentage of all particles produced by transfected 5B8 cells that are vesicles and GFP-loaded ARMMs, we performed a single-particle analysis of the samples using Single particle analysis (Figure 3). The size histogram suggested that the 5B8 cell line produced particles with two different size distributions, with one peak at ∼50 nm and the other at ∼ 80 nm. ARMMs (i.e., GFP^+^ events) were more frequently located within the smaller peak (Figure 3A and B). Approximately 50% of the particles were GFP^+^, which was in agreement with the NTA data showing a ∼2-fold increase in particle production when ARRDC1 was overexpressed.

**FIGURE 3:**
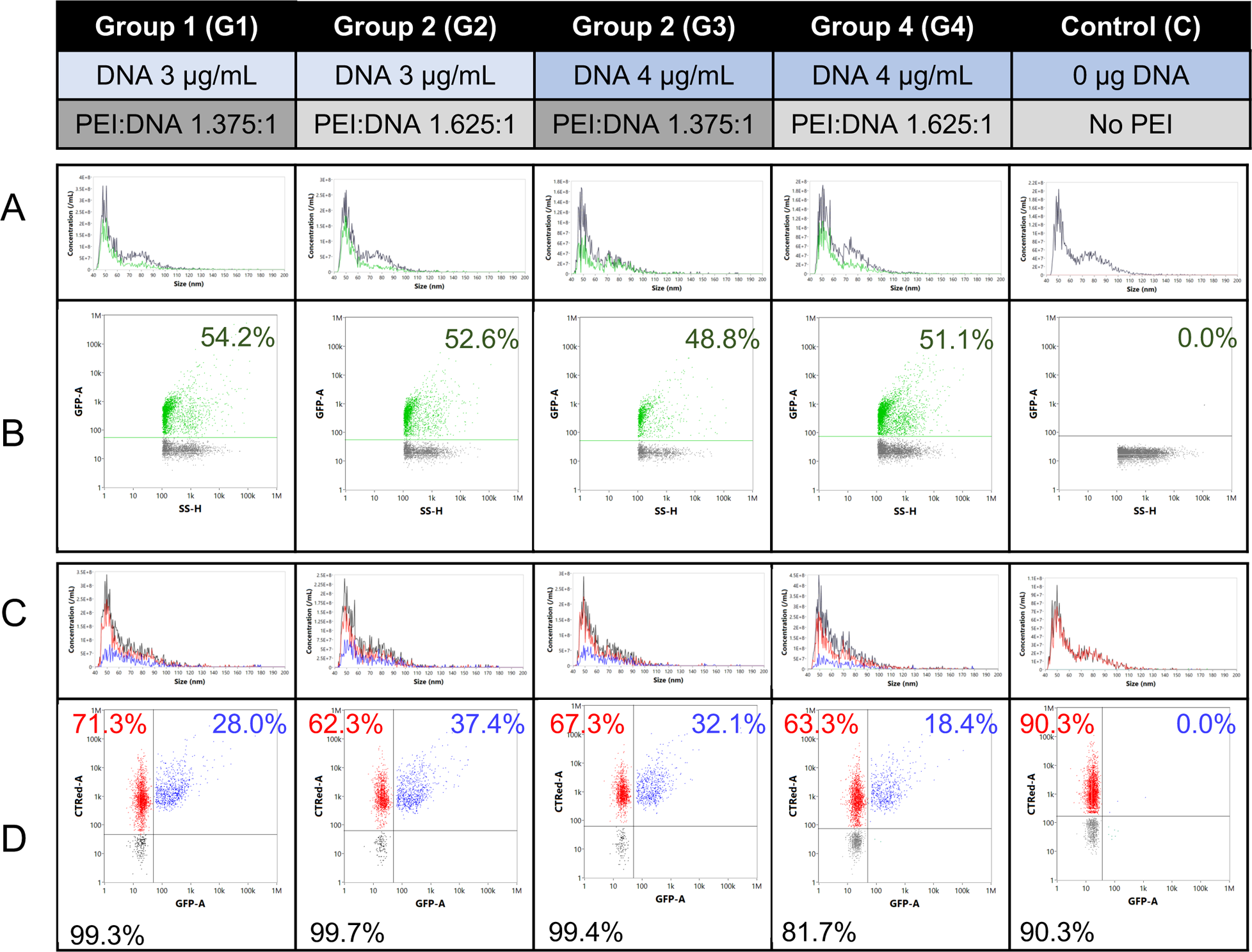
Single particle analysis analysis to characterize the GFP^+^ particles as intact vesicles. **(A, B)** Size histograms and GFP^+^ events (green dots on dot plot) detected in the raw conditioned medium. **(C, D)** After labeling and wash-out of free dye, the % positive for CTRed (red), GFP (green), and double-positive (blue) was determined. The total CTRed^+^ population % is denoted in black.

Next, we labeled the intact vesicles from G1–4 with CTRed, and the free dye was removed using SEC. Most particles (> 99%) derived from G1–3 were CTRed^+^, indicating that these were intact vesicles. Notably, there were no GFP^+^/CTRed^−^ events (Figure 3D), indicating that no damaged particles or aggregates containing GFP were detected. The presence of CTRed dye may overwhelm the GFP signal of weakly positive/very small ARMMs, causing the apparent % GFP^+^ (Figure 3D) to decrease when double-stained (Figure 3D) compared to single staining (Figure 3B). We also labeled the particles with antibodies to tetraspanins to determine whether these are a useful biomarker of ARMMs, and found partial co-localization (Supplemental Figure 2).

### 3.2 Engineering a stable 5B8 producer cell line to support scalable production of ARMMs

We transduced 5B8 cells with lentiviruses to generate cells stably expressing ARRDC1-GFP. After lentiviral transduction and selection, the expression of ARRDC1-GFP was confirmed by imaging of the cells and evaluating the cell lysates and ARMMs by Western blotting (Figure 4A-B). We also performed a flow cytometric analysis of the stable cell line, which showed that they were approximately 79% GFP^+^ (Figure 4E). We evaluated these cells for the production of GFP-loaded ARMMs.

**FIGURE 4:**
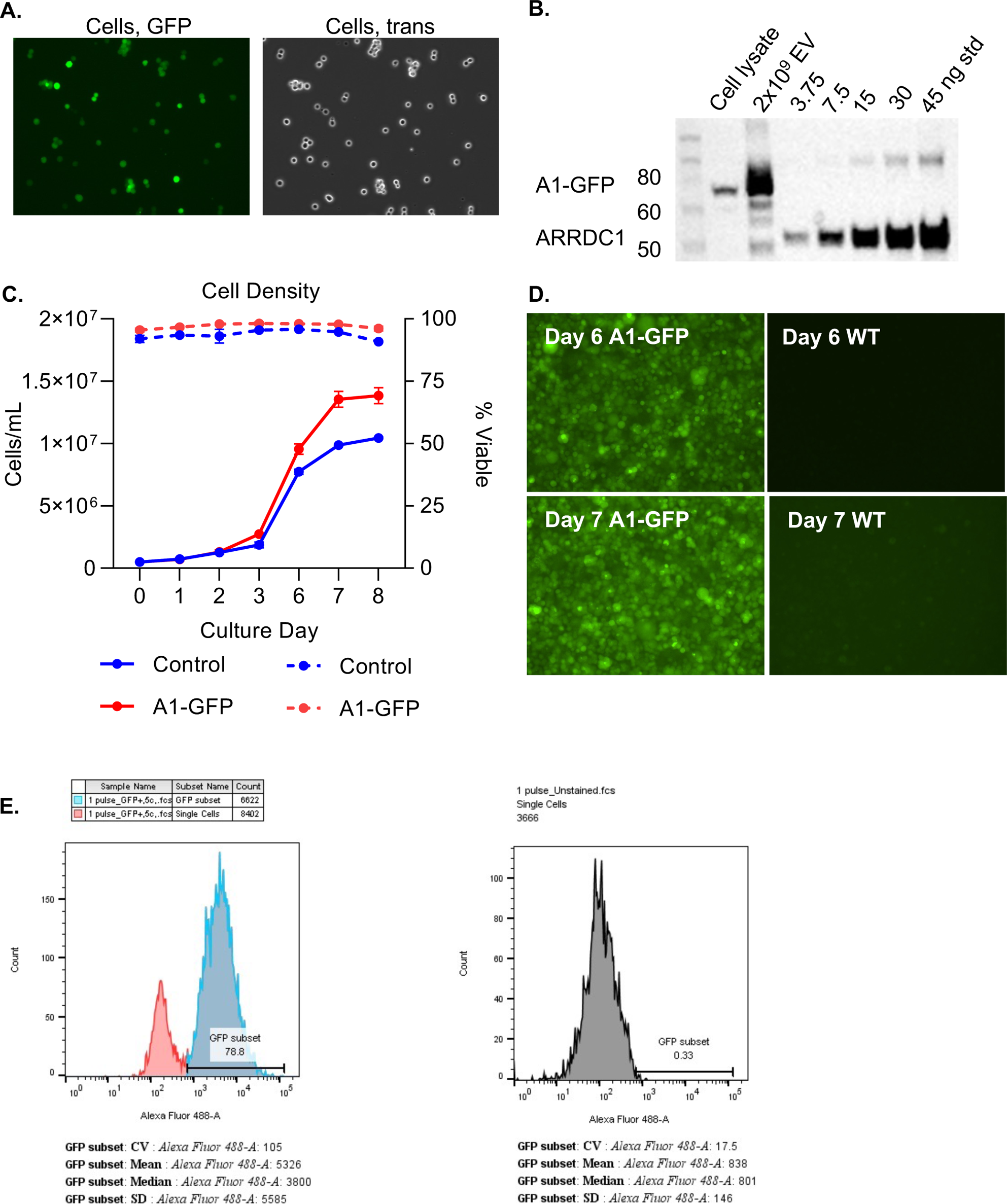
**(A)** The 5B8 cells were almost all GFP^+^ after selection. **(B)** A1-GFP was significantly expressed in the cell lysate and EVs from the stable line. **(C)** Viable cell density (solid lines) and % viable (dashed lines) each day after seeding the cells in shake flasks. **(D)** Representative images of the cells on days 6 and 7 after seeding showed maintained expression of A1-GFP and high density. WT cells on the same days are shown for comparison, with only mild autofluorescence visible. **(E)** Flow cytometry of the WT and transformed cells is shown as histograms of 488 (GFP) intensity.

The stable cell line was cultured in shake flasks to evaluate seeding density, cell growth, maximum VCD, and ARMM production before being used to produce CM in a 3 L stirred tank bioreactor. Control and ARRDC1-GFP stable 5B8 cells were cultured in 1 L shake flasks to produce 300 mL of CM. Two seeding densities were tested, 0.5 and 1 × 10^6^/mL. We observed that a 0.5 × 10^6^/mL seeding was sufficient to allow for steady growth and high viability for 7 d in culture. By the 8^th^ day, the cell number plateaued, and viability began to decrease. Expression of the fusion protein did not adversely impact cell growth; the ARRDC1-GFP stable line grew to a slightly greater density than the control line (1.4 × 10^7^ vs. 1 × 10^7^) and had a slightly higher VCD. Both groups showed > 90% viability throughout the experiment (Figure 4C-D).

Single particle analysis showed a concentration of 1.82 × 10^10^ particles/mL in the CM of the control cells, a median size of ∼55 nm, and an average of 61 nm (Figure 5A-B). In the A1-GFP CM, the concentration remained constant at 1.82 × 10^10^ particles/mL, and there was a bi-phasic distribution of sizes as observed in transfected cells. The particles were 55.6% GFP^+^, similar to the transfected cells.

**FIGURE 5:**
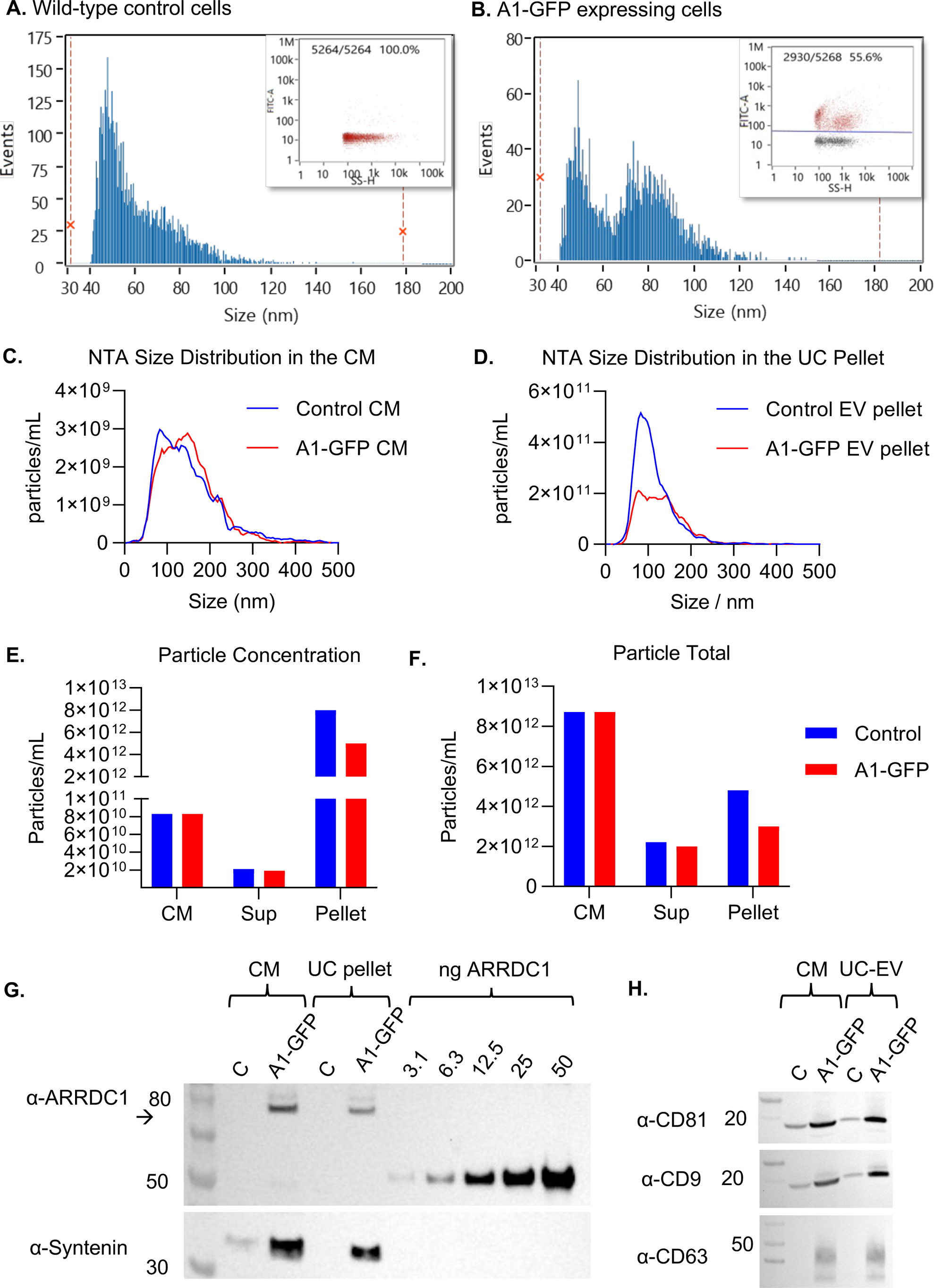
Single particle analysis data on particles from the CM of **(A)** the WT and **(B)** the A1-GFP-expressing cells. Insets show the dot plot of green fluorescence intensity vs. side scatter. **(C–D)** NTA-generated size distributions for particles in the conditioned media and UC pellet. Note that the bi-phasic peak is seen in the A1-GFP EVs but not the control EVs by NTA as well as Single particle analysis. **(C)** NTA particle concentrations in the CM, Sup, and Pellet. **(F)** Mass balance of total particles. **(G)** Western blot for ARRDC1 and Syntenin on 1 × 10^9^ particles from the CM and UC pellet from the control and A1-GFP stable line, including a standard curve of known recombinant ARRDC1 (ng). **(H)** Western blot for tetraspanins CD81, 9, and 63 in the control and A1-GFP CM and UC pellet.

Next, 105 mL of each CM were ultracentrifuged, and the EVs were resuspended in 600 µL of PBS. After determining the particle concentration by NTA, CM- and UC-pelleted particles were tested by Western blot analysis, loading 1 × 10^9^ particles/lane. Interestingly, the NTA size distribution also appeared to be slightly bi-phasic in the A1-GFP UC-EVs, although it was less distinct than that provided by Single particle analysis (Figure 5C-F). As before, the A1-GFP band was more intense in the CM than in the pelleted EVs, (Figure 5G). CD9, CD63, and CD81 were all detected in the A1-GFP particles (Figure 5H). Using the standard curve generated with ARRDC1 purified protein, we determined the amount of ARRDC1 immunopositive signal in the control and A1-GFP EVs and determined the mass and number of molecules of A1-GFP per particle. ∼76 ± 10 molecules of ARRDC1/EV were calculated from pelleted EVs (Table 2).

**TABLE 2:** Determination of the number of molecules of ARRDC1/EV by Western blot with a standard curve.

| | Pixel Density (AU) | ng in lane | Volume loaded (μL) | μg/mL | particles/mL | Molecules/particle |
| --- | --- | --- | --- | --- | --- | --- |
| A1-GFP CM | 1149 | 7.03 | 12.05 | 0.58 | 8.3e10 | 92 |
| A1-GFP UC-EV | 576 | 4.96 | 0.2 | 24.78 | 5.0e12 | 65 |

### 3.3 Production of ARMMs using a scalable stirred tank bioreactor and downstream processes

To establish a seed train, cells stably overexpressing ARRDC1-GFP were first expanded into a 1-L shake flask and seeded at 0.5 × 10^6^/mL in 300 mL. This culture was used to seed the 3 L bioreactor once it reached ∼5 × 10^6^ viable cells/mL (Figure 6A). Two additional shake flasks (300 mL) were used to monitor cell growth and fluorescence during the 7-day expansion. After seeding in the bioreactor, the cells were cultured for six days (Figure 6B).

**FIGURE 6:**
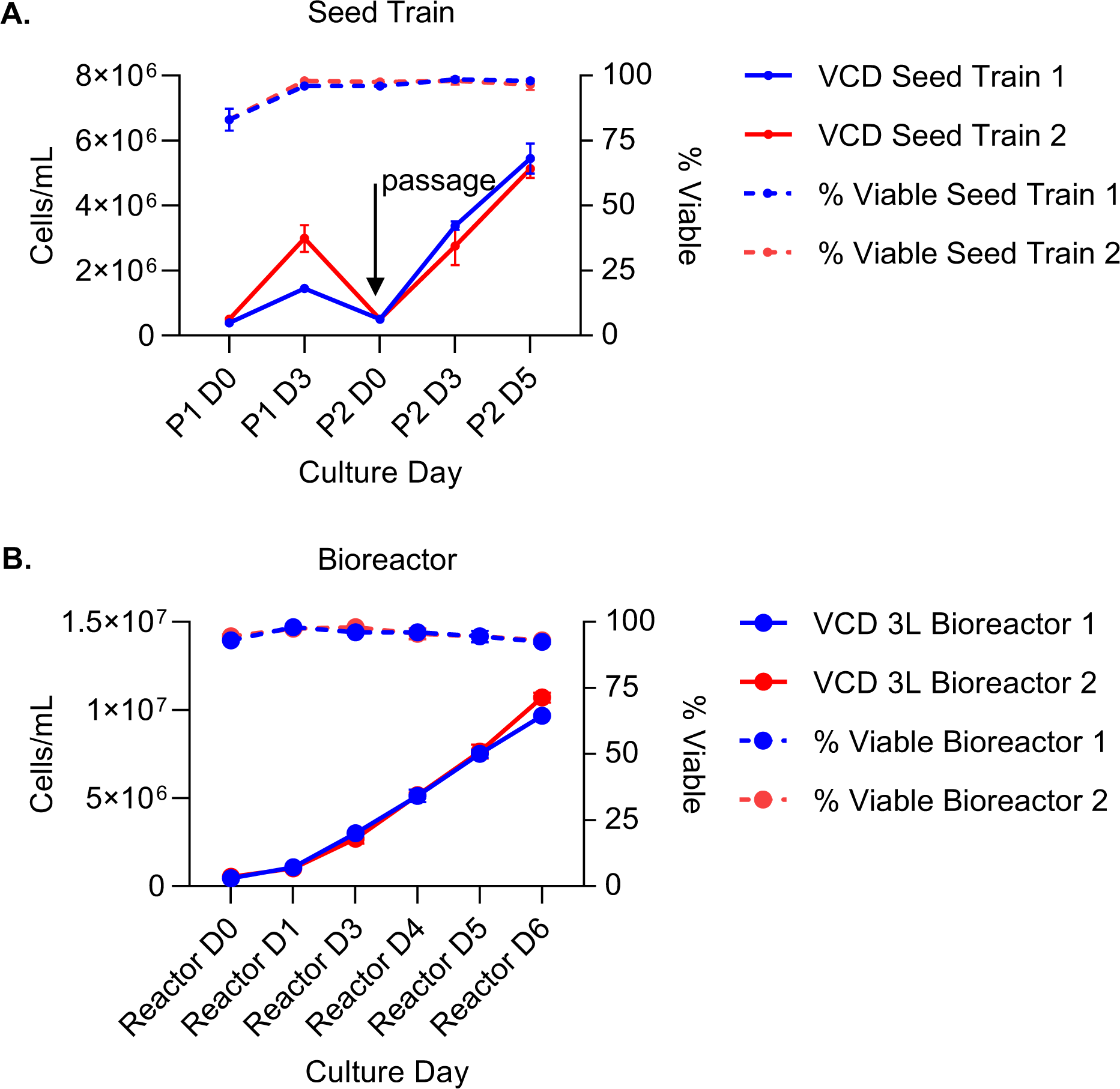
Cell expansion from seed train to stirred tank bioreactor. **(A)** Cells were seeded (day 0) and then passaged 3 and 5 days later (n = 2 replicates in two separate experiments). Cell density (solid lines) and viability (dashed lines) were comparable. **(B)** Cell density (solid lines) and viability (dashed lines) were similar, and viability was very high throughout the upstream process.

#### Downstream process of ARMM purification

CM was sampled on day 6 for cell count and viability and then harvested by removing cells and large debris using depth and 0.2 µm sterile filtration (i.e., clarified CM). The clarified CM was maintained overnight at 4 °C. A total of 550 mL was removed for sampling to obtain a UC sample as a control. To separate EVs from host cell proteins by size and to concentrate samples, TFF was conducted, followed by buffer exchange to prepare for AEX. The retentate was 0.45 µm filtered (TFF Isolate) to eliminate debris and large aggregates. The remaining volume (197 mL) was resolved in an AEX column, and the column was washed with 80 mL of buffer. Fractions were eluted using a linear salt gradient, and those containing EVs were pooled to a final volume of 80 mL, referred to as the AEX eluate. The AEX eluate was further purified using multimodal chromatography. This was buffer-exchanged with a final TFF and sterile filtered. Table 3 lists sample names, processing steps, and total volumes.

**TABLE 3:** Downstream processing operations, sample naming, and volumes.

| **Test Article** | **Description** | **Total volume represented by sample (mL)** |
| --- | --- | --- |
| Clarified CM | DNAse treatment, salt added, bioreactor flush, depth filtered, sterile filtered | 4463 |
| TFF Input | Same as above but with 550 mL removed for UC/sampling | 3919 |
| TFF Isolate | TFF1-diafiltered concentrate | 197 |
| AEX Eluate | Chromatography-pooled fractions | 80 |
| Final purified EVs | Final sterile filtration after TFF2 | 73 |

The particles were retained by TFF, resulting in a 20x concentration (Figure 7A and B, and Table 4). The total amount of GFP decreased during this step, suggesting the wash-out of some non-vesicular GFP. There was a further 2.5 × volume reduction by AEX, which was associated with some loss of particles and GFP; however, the second TFF step, resulting in the final purified EVs, showed no further reductions. TFF efficiently removed the host cell proteins, as measured by the Bradford assay, by 86%, and AEX further reduced this to only 5% of the input. The second TFF step resulted in a slight reduction in protein levels (Figure 7C). The enrichment of particles per microgram of protein was highest after the first TFF (Figure 7D). A second 3 L bioreactor run yielded similar results to the first; the particle concentration and per-step mass balance are shown (Figure 7E and F).

**FIGURE 7:**
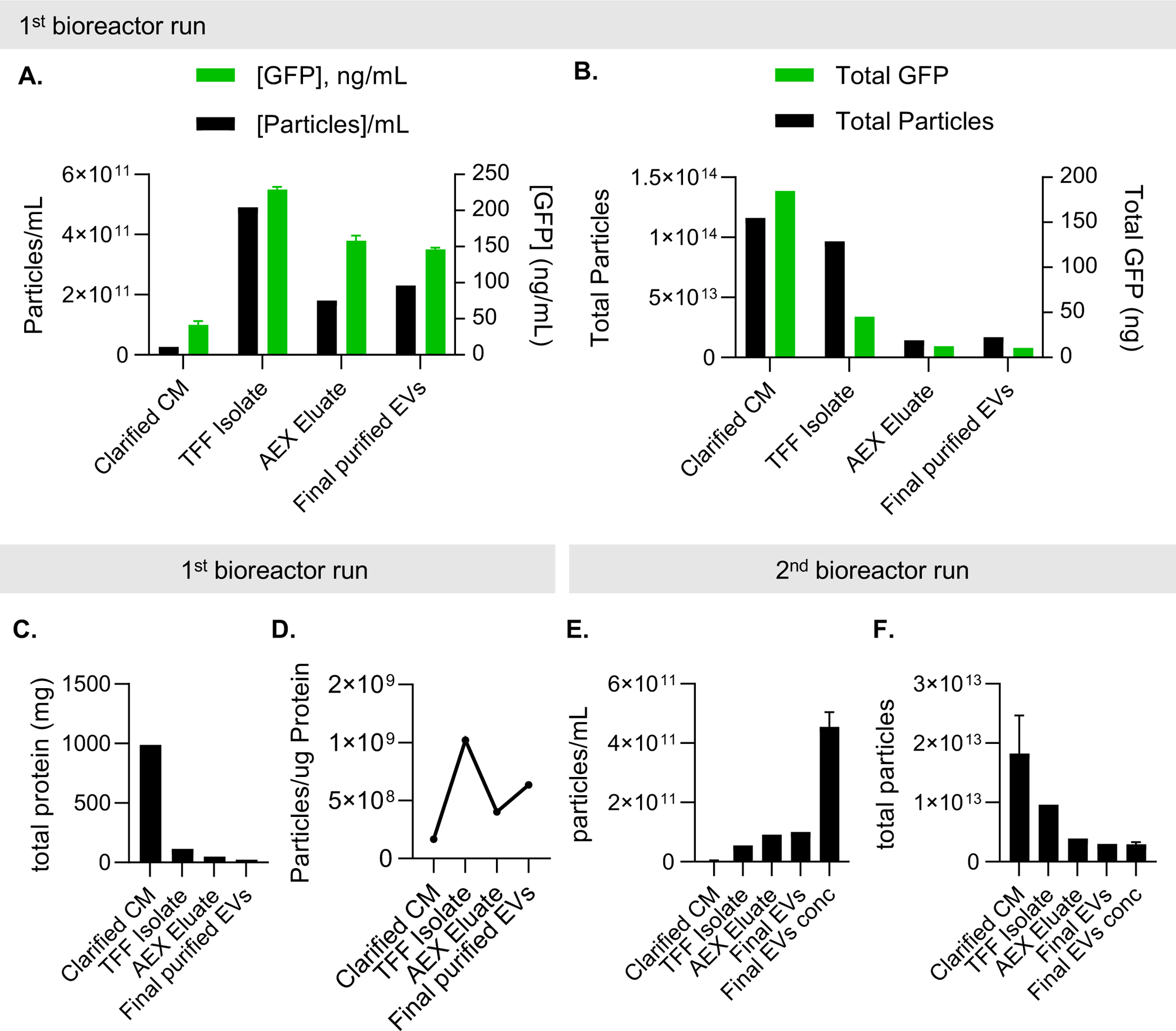
Characterization of the particles throughout the downstream process **(A)** Concentration of particles was determined by NTA, and GFP concentration was determined by ELISA at each downstream processing step. **(B)** The mass balance of particles and GFP, concentration x volume; **(C)** the total amount of protein was determined by Bradford protein assay. **(D)** The ratio of particles per μg total protein for each step. **(E)** Particle concentration and **(F)** total particles for the second bioreactor run were similar to the first run, but the final purified EVs were further concentrated to prepare them for use *in vivo*.

**TABLE 4:** Mass balance of particles, ARRDC1, and GFP per downstream step

| | Vol<br>(mL) | [particles]/mL | Total<br>Particles | Particle<br>Yield | GFP<br>ELISA<br>(ng/mL) | Total<br>GFP<br>(μg) | GFP<br>Yield | Protein<br>(μg/mL,<br>Bradford) | Total<br>protein<br>(mg) | Protein<br>Yield |
| --- | --- | --- | --- | --- | --- | --- | --- | --- | --- | --- |
| TFF Input<br>(Clarified<br>CM) | 3919 | 2.60E+10 | 1.02E+14 | 100% | 41.5 | 163 | 100% | 175.4 | 687 | 100% |
| TFF<br>Isolate | 197 | 4.90E+11 | 9.65E+13 | 95% | 228.8 | 45 | 28% | 479.9 | 95 | 14% |
| AEX<br>Eluate | 80 | 1.80E+11 | 1.44E+13 | 14% | 158.1 | 13 | 8% | 448.9 | 36 | 5% |
| Final<br>purified<br>EVs | 73 | 2.30E+11 | 1.68E+13 | 16% | 145.9 | 11 | 7% | 361.6 | 26 | 4% |

We tracked the expression of ARMMs-associated proteins retained at each step by loading 5 × 10^8^ particles per lane for Western blot. The cell lysate was included as a control; this was the only lane that showed the presence of Calnexin, suggesting that there was no contamination with cellular vesicles (Figure 8A). Evaluation of ARMMs loading efficiency using the standard curve revealed that there were approximately 112 ARRDC1-GFP molecules/EV in the final product. CD63, CD9, and CD81 were present (Figure 2E and 5H) and remained relatively consistent throughout the process (Figure 8B). Single particle analysis was used to determine the size and GFP positivity of the samples obtained at each process step (Figure 8C-D). The two size peaks were visible in the TFF Isolate and AEX eluate but appeared less distinct in the final EVs and final EV concentrated samples. As mentioned previously, the peak of the GFP^+^ population tended to be located among the smallest particles in the histogram. GFP positivity was increased after the first TFF, likely due to the removal of proteins and other contaminants from the CM. Then, it remained ∼40 ± 10% through the rest of the process (Figure 8C–D). CTRed labeling was performed on a second lot of purified ARMMs (2^nd^ bioreactor run), showing that 99% of the particles were positive; that is, the GFP^+^ particles were likely to be intact ARMMs.

**FIGURE 8:**
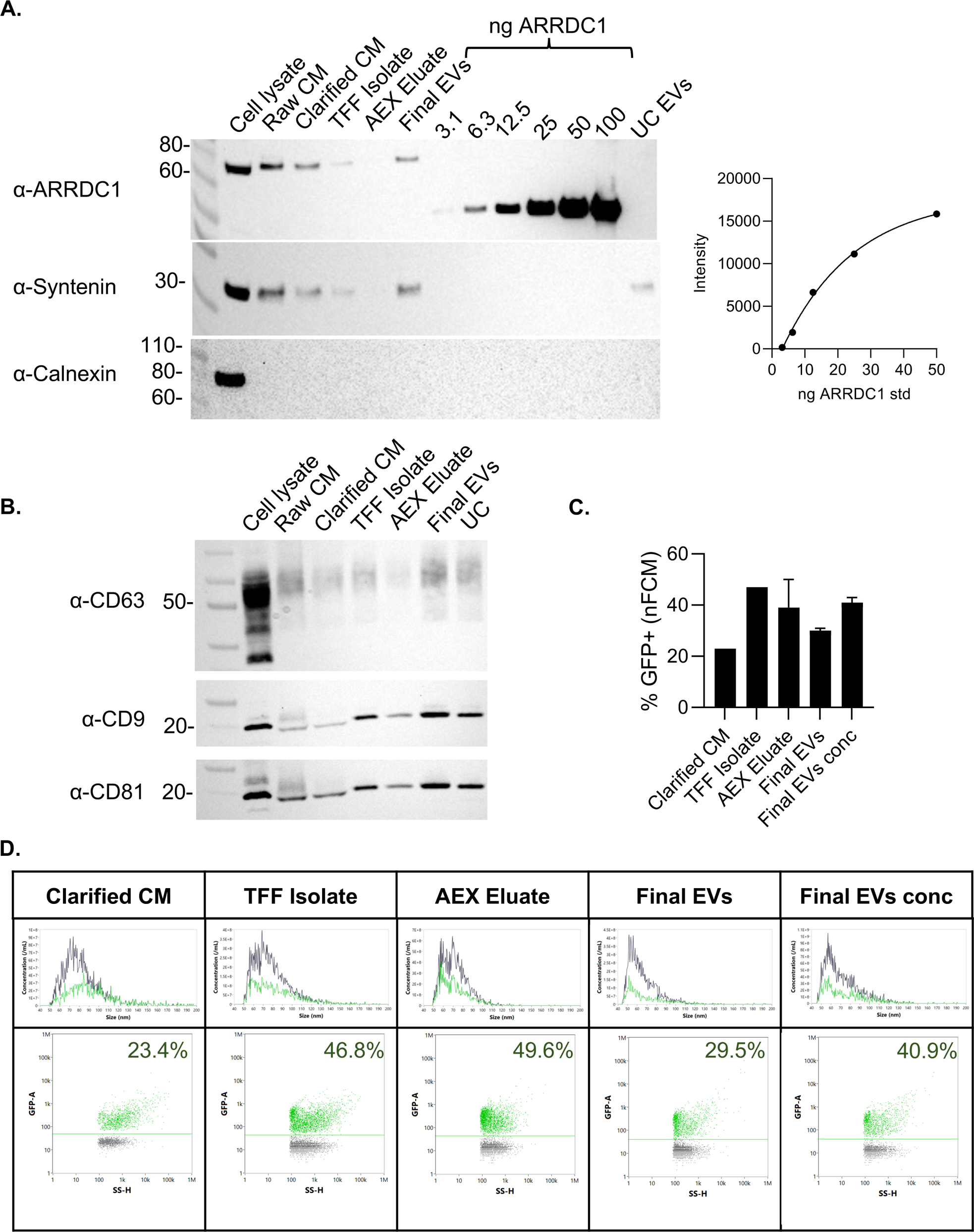
**(A)** Western blot on the cell lysates and the isolated ARMM particles during different downstream processing steps. A standard curve of recombinant ARRDC1 was used to determine the amount of ARRDC1 in the samples. **(B)** Western blot was performed under non-reducing conditions to assess the expression of tetraspanins. **(C)** Single particle analysis was used to determine % of particles that were GFP^+^. **(D)** Representative size distribution curves and dot plots from nFCM analysis. The green population indicates the GFP^+^ events vs. side scatter for each unit of operation taken into account in bar graph C.

### 3.4 Bioactivity, biodistribution, and pharmacokinetics of 5B8 ARMMs

In parallel with the scalable downstream process described above, 200 mL of clarified CM from the second 3 L bioreactor run was purified by ultracentrifugation, and the uptake of the resulting ARMMs was compared in A549 cells. The final EVs from the TFF/AEX/TFF process had higher uptake in A549 cells than UC-EVs (Figure 9A), suggesting that the purification steps did not cause any damage that would prevent them from interacting with the cell surface or decrease potential bioactivity.

**Figure 9:**
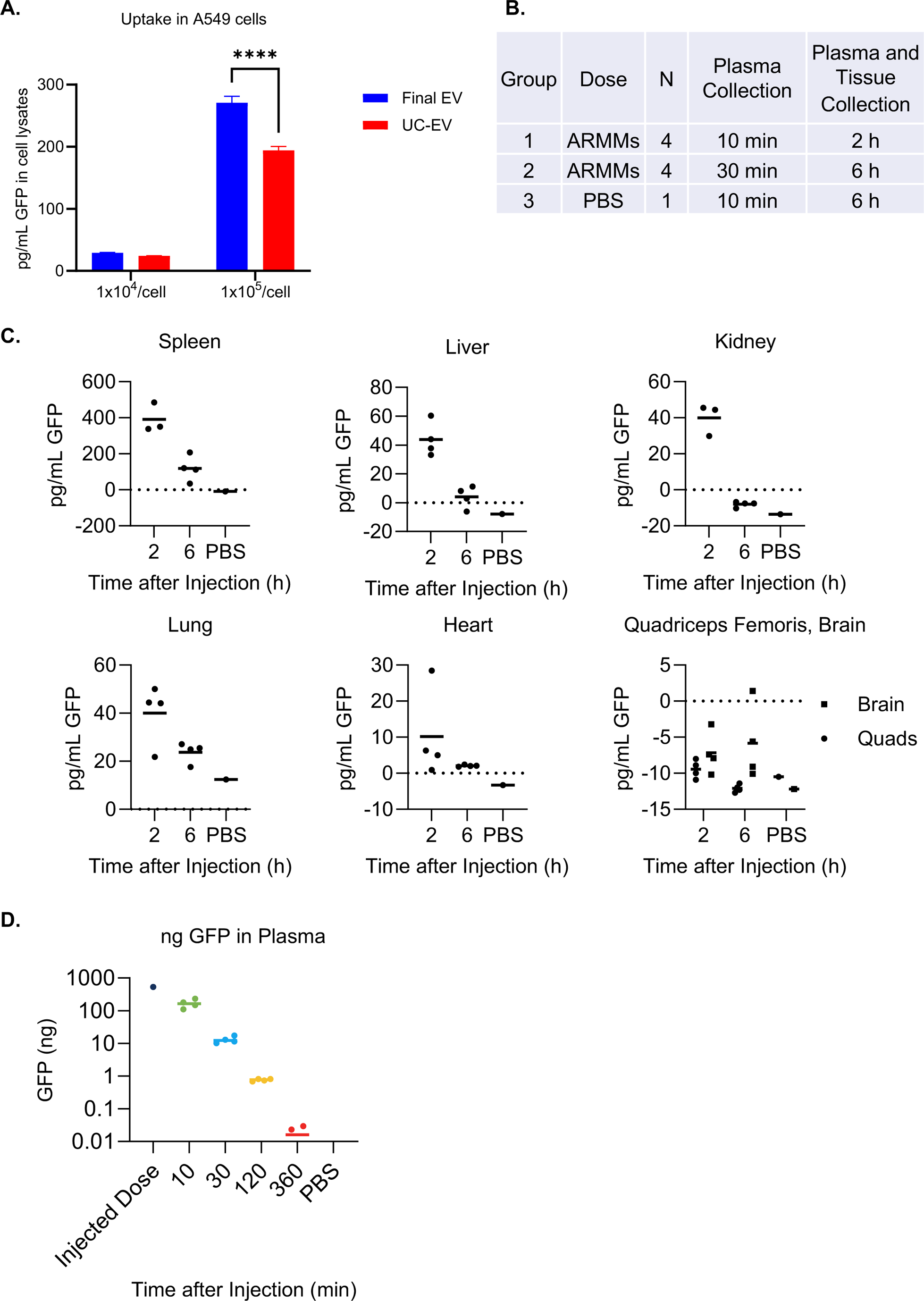
**(A)** A549 cells were treated with two different doses of ARMMs per cell for 24 h, and GFP concentration in the cell lysates was assessed by ELISA. ARMMs processed using TFF and AEX showed higher uptake than those processed by UC (*P* < 0.001, n = 3). **(B)** Mice were assigned to groups to receive PBS or ARMMs (GFP) with tissue collection at 2 h or 6 h, each with an earlier plasma collection timepoint. **(C)** GFP concentration in mouse tissue homogenates was assessed by ELISA, with 300 μg total protein per well. **(D)** GFP concentration was assessed in plasma over time by ELISA.

Next, *in vivo* biodistribution and pharmacokinetics were assessed in mice after intravenous (IV) injection. The study timeline is presented in Figure 9B. The concentration of GFP in tissue homogenates 2 and 6 h after injection showed that ARMMs rapidly accumulated in the spleen and, to a lesser extent, in the liver, kidneys, and lungs, after which they gradually broke down and remained detectable by the 6-h timepoint (Figure 9C). 5B8-derived ARMMs demonstrated a plasma half-life of 6 ± 0.4 minutes (Figure 9D).

## 4 Discussion

The ability to produce ARMMs, and more broadly EVs, using scalable processes has implications for their potential use in therapeutics. Various types of producer cells and processes have been used to generate EVs with diverse properties[26]. These include a range of adherent and suspension cell types, as well as stem and primary cell-derived EVs [27,28]. Although benchtop studies on EVs produced by such systems are useful for evaluating their molecular characteristics and beneficial properties, scalable production remains a major hindrance [27]. To enable therapeutic use of acellular products, such as ARMMs, it is necessary to establish a framework for large-scale production of EVs. Ideally, ARMMs production would be carried out in a suspension cell line that can be grown to high densities in an FBS-free medium lacking animal products and would produce sufficient amounts to support *in vivo* evaluation.

Previous efforts to generate ARMMs had relied on transfection-based methods in a 293T adherent cell line [10,13]. In this study, we evaluated EV productivity of the HEK293-derived suspension cell line 5B8 developed by Lonza using transient transfection or by stable expression of a cassette encoding ARRDC1-tethered GFP. ARMMs produced using either approach yielded efficient GFP loading at high percentages (48-55%). Production of ARMMs using the 5B8-derived stable producer cell line demonstrated the potential of this cell line to support scaled production to enable large animal studies and, importantly, future evaluation in the clinic. Particular attention was paid to the viability of producer cells to attempt to limit levels of contaminants during ARMMs purification. In general, cell viability levels were maintained at >80%. Using a stable producer cell line approach provided a significant advantage in this regard resulting in a VCD of ∼ 90% during harvesting of ARMMs-containing CM. Several approaches to scale production have been previously discussed [27,29]. To our knowledge, this is the first demonstration of scaled production of engineered EVs using a stable cell line producer system and in the absence of inducers or stressors, which could potentially compromise the quality of the ARMMs product.

A range of extracellular vesicle purification strategies have been evaluated or proposed by others [30]. However, most bench scale production efforts continue to rely on precipitation or UC methods [31]. Here, we used two approaches to isolate ARMMs. As noted, ultracentrifugation is a convenient and commonly used approach for small-scale purification of EVs, including ARMMs; however, it has limitations when used to process large liter-scale CM to purify ARMMs. We used a combination of ultrafiltration and chromatography-based approaches to isolate ARMMs from larger volumes of CM. The characterization of ARMMs using either approach generated comparable product recovery rates with more favorable product characteristics, including improved loading and uptake, by chromatography. Importantly, ARMMs produced by UC, or the downstream processing approach delineated herein, were active and showed robust uptake *in vitro*.

To interrogate ARMMs biodistribution, we intravenously administered GFP-loaded ARMMs to mice and investigated their uptake across a large range of tissues. A strong signal was observed in the spleen and liver and, to a lesser extent, in the kidney, supporting the potential use of ARMMs in the delivery of therapeutic payloads to these target tissues. Other studies have reported similar biodistribution of EVs in highly vascularized organs [32], suggesting that the methods for production and purification utilized here do not alter the properties of ARMMs. The pharmacokinetics and half-life were also similar to that previously described for other types of EVs administered intravenously [33,34]. Of note is the potential impact of dose on biodistribution, which was not evaluated in this study.

Altogether, this work illustrates a scalable framework for production of engineered ARMMs using Lonza’s suspension cell line 5B8. While multiple methods are currently used for purification of EVs, therapeutic evaluation will require a streamlined and scalable purification process, such as the chromatography-based approach described here, to enable reproducible generation of high-quality product. Such processes will permit evaluation of a range of therapeutic strategies using ARMMs as modular non-viral vehicles for delivery of Cas/gRNA complexes, proteins, and RNA.

## 5 Acknowledgments

The graphical abstract was created with Biorender.com.

## Data availability statement

All data generated or analyzed during this study are included in this published article and its supplementary information files.

## Ethics approval statement

The animal research was approved by the Institutional Animal Care & Use Committee at the Charles River Accelerator and Development Lab.

## Appendices

**Supplemental Figure 1.**
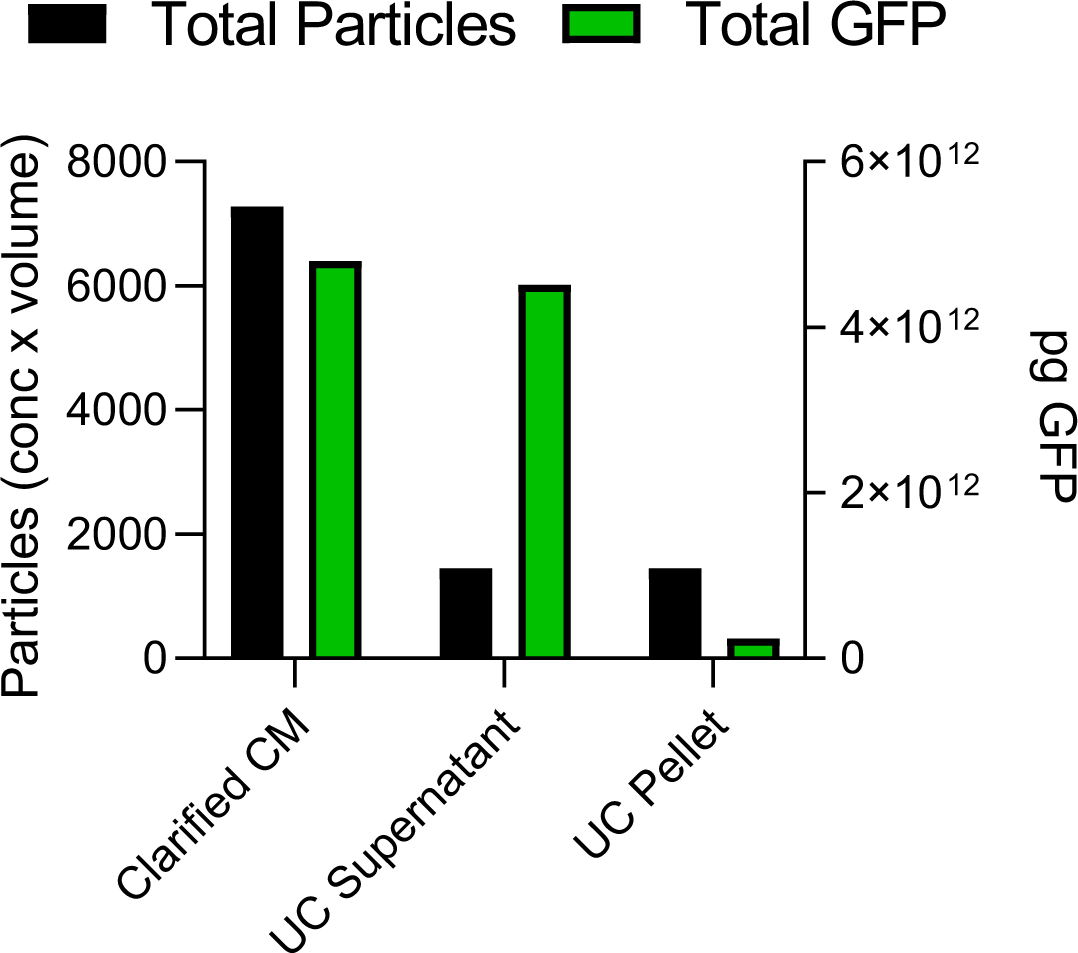
Distribution of particles and GFP after UC. Particle concentrations were determined by NTA, and GFP concentrations were determined by ELISA. The total amount (concentration x volume) of each is shown in the CM, supernatant, and re-suspended pellet.

**Supplemental Figure 2.**
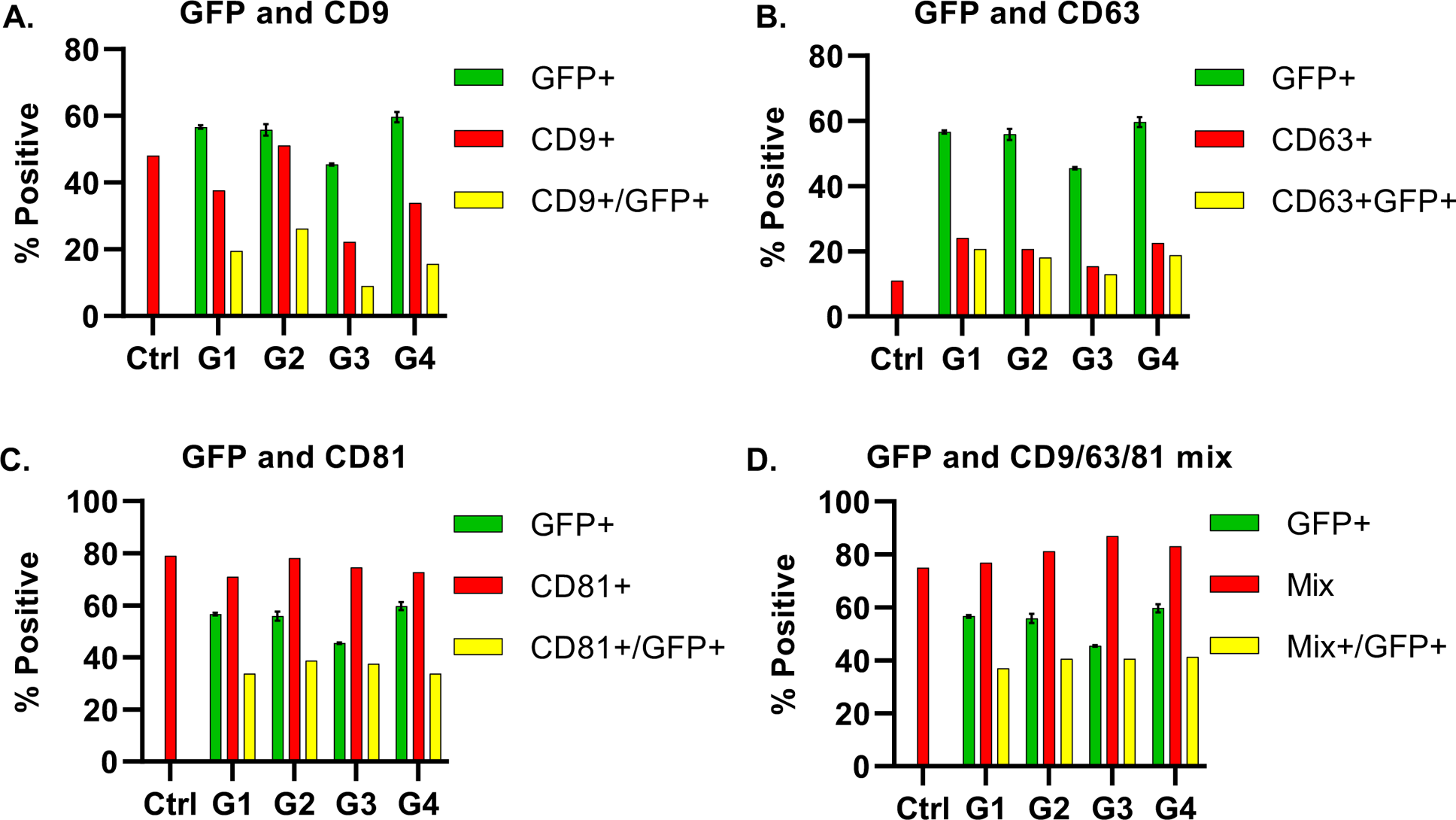
GFP co-localization with tetraspanins. Particles from the control and transfected samples were labeled with antibodies to **(A)** CD9, **(B)** CD63, **(C)** CD81, and **(D)** a mix of these, and analyzed by single particle analysis. For each chart, the red bar indicates the percentage of events that were labeled with the antibody, and the green bar represents the percentage of GFP+ events. The yellow bar indicates the events that were positive for both GFP and the antibody label.

---

- The engineering of ARRDC1-mediated microvesicles (ARMMs) as non-viral gene therapy vehicles could overcome challenges associated with other delivery approaches.
- Our study explored scalable strategies for generating GFP-loaded ARMMs from Lonza’s suspension HEK293 cells.
- Downstream purification utilized Tangential Flow Filtration (TFF) and Anion Exchange Chromatography (AEX).
- ARMMs were characterized by single particle analysis, nano-tracking analysis, immunoblotting, and ELISA.
- *In vivo* studies in mice revealed that purified ARMMs are active and showed biodistribution of ARMMs to the spleen, liver, kidneys, and lungs, highlighting their potential for therapeutic delivery.

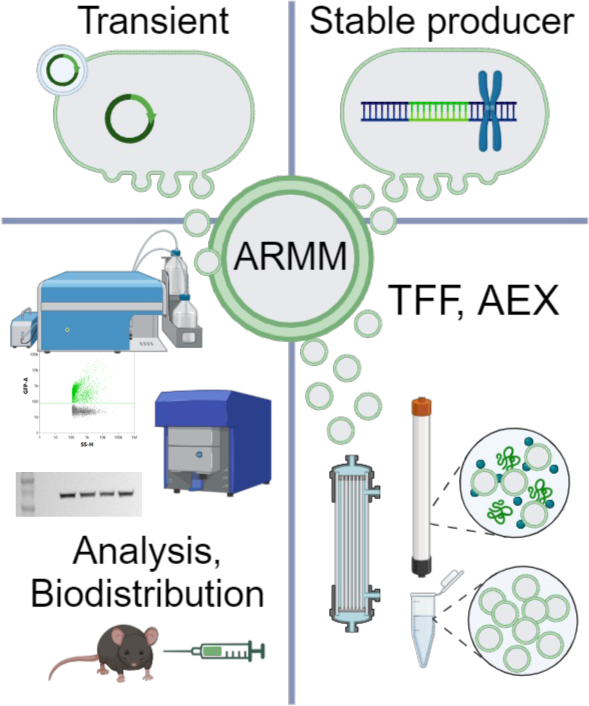

